# Simultaneous enumeration of yeast and bacterial cells in the context of industrial bioprocesses

**DOI:** 10.1101/2024.06.26.600471

**Authors:** Carolina Teixeira Martins, Ana Paula Jacobus, Renilson Conceição, Douglas Fernandes. Barbin, Helena Bolini, Andreas Karoly Gombert

## Abstract

In scenarios where yeast and bacterial cells coexist, e.g. the food and bioethanol industries, it would be of interest to simultaneously quantify the concentrations of both cell types, since traditional methods used to determine these concentrations individually take more time and resources. Here, we compared different methods for quantifying the fuel ethanol *Saccharomyces cerevisiae* PE-2 yeast strain and cells from the probiotic *Lactiplantibacillus plantarum* strain in microbial suspensions. Individual suspensions were prepared (∼10^7^, yeast cells/mL or ∼10^9^, bacterial cells/mL) and mixed in 1:1 or 100:1 yeast-to-bacteria ratios, covering the range typically encountered in sugarcane biorefineries. The following methods were used: bright field microscopy, manual and automatic Spread-plate and Drop-plate counting, flow cytometry (at 1:1 and 100:1 ratios), and a Coulter counter (at 1:1 and 100:1 ratios). By subjecting the same mixed cell suspension to each technique, we observed that for yeast cell counts in the mixture (1:1 and 100:1 ratios), flow cytometry, the Coulter counter, and both Spread-plate options (manual and automatic CFU counting) yielded statistically similar results, while the Drop-plate and microscopy-based methods gave statistically different results. For bacterial cell quantification, the microscopy-based method, Drop-plate, and both Spread-plate plating options and flow cytometry (1:1 ratio) produced no significantly different results (P>0.05). In contrast, the Coulter counter (1:1 ratio) and flow cytometry (100:1 ratio) presented results statistically different (P<0.05). Additionally, quantifying bacterial cells in a mixed suspension at a 100:1 ratio wasn’t possible due to an observed overlap between yeast cell debris and bacterial cells. The results from this work indicate that each method has limitations, advantages, and disadvantages, meaning that the best option will always depend on the application. We present a comparison of the techniques, in terms of time-to-results, cost of analysis and equipment, range of detectable cell/particle diameters, adequacy for simultaneous enumeration, and general pros and cons.

**Graphical Abstract:** This study compares methods for simultaneously quantifying yeast and bacterial cells in a mixed sample, highlighting that in different cell proportions, some methods cannot quantify both cell types and present distinct advantages and limitations regarding time, cost, and precision.

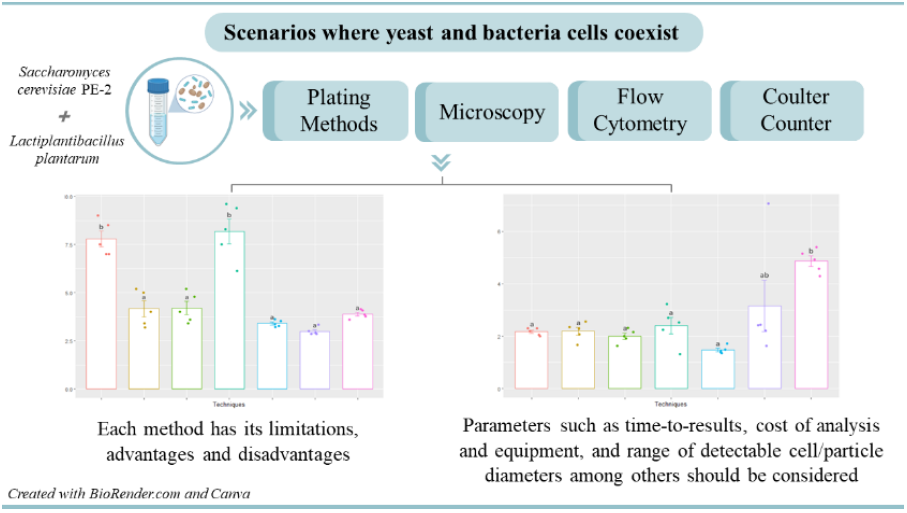

## INTRODUCTION

The enumeration of yeast and bacterial cells is a critical task in diverse scientific and industrial domains, encompassing microbiology, biotechnology, food, medicine, and environmental studies (Trinh & Lee, 2022; Vinderola et al., 2019). These microorganisms play pivotal roles in fermentation, food production, wastewater treatment, and the pathogenesis of infectious diseases (Doppler et al., 2022; Trinh & Lee, 2022; Vinderola et al., 2019). Precise measurement of yeast and bacterial concentrations is essential for comprehending, monitoring, and controlling biotechnological processes, as well as for disease diagnosis, and water and food monitoring, among other applications (Beckner et al., 2011; Wang et al., 2021). For instance, in the food and beverage industry, it is vital to maintain consistency in fermented products, such as bread, be er, and probiotic-rich beverages like kefir (Garofalo et al., 2015; Williamson et al., 2023). In biorefineries, quantification is crucial for monitoring fermentation processes and controlling contaminating bacterial cells (Ceccato-Antonini, 2012., Ceccato-Antonini 2018). The pharmaceutical industry relies on precise quantification to optimize yields and guarantee the safety and efficacy of drugs and vaccines produced through bioprocessing (Deshmukh et al., 2016; Fontana et al., 2017). In environmental applications, quantifying microbial populations is essential for assessing wastewater treatment efficiency and environmental bioremediation efforts (Douterelo et al., 2014; Lee et al., 2021). Additionally, in medical research and clinical diagnostics, quantifying yeast and bacterial cells is fundamental for understanding microbiome dynamics, studying infections, and monitoring treatment effectiveness. Overall, accurate quantification of both yeast and bacterial cells in scenarios where they coexist is vital in diverse sectors, ensuring the success of processes, the quality of products, and the integrity of environmental and human health.

Within diverse fields employing yeast and bacterial cells, the selection of microbial enumeration methods varies. The evaluation and selection of these methods are a crucial step in determining which technique is most appropriate to meet the specific needs o f the study or application in question. Some parameters to consider for this choice are: i) Sensitivity and detection range; ii) specificity; iii) time and speed; iv) complexity and training; v) cost; vi) identification capacity; vii) accuracy; viii) detection of live and dead cells; ix) generated environmental impact;x) operator interference; and xi) sample complexity. (Douterelo et al., 2014; Vinderola et al., 2019) In medical and disease diagnostics, prevalent techniques for pathogen detection and quantification encompass culture-based techniques like agar plate counting with suitable growth media and nucleic acid-based methods such as PCR, qPCR, and amplicon sequencing (Trinh & Lee, 2022). In the food sector, especially in probiotic development, techniques like traditional plating, optical microscopy, nucleic acid-based methods (PCR and RT-PCR), direct imaging methods, and flow cytometry are frequently employed for cell enumeration (Davis, 2014; Vinderola et al., 2019). In biorefineries, where *Saccharomyces cerevisiae* is of primary interest, methods such as direct counting through bright-field microscopy and classic serial dilution and plating on appropriate culture media are employed to monitor contaminating bacteria (Beckner et al., 2011; Ceccato-Antonini, 2012; Vieira & Fernandes, 2012).

The conventional spread plating method, commonly performed using Petri dishes, is extensively used in microbiological analyses due to its cost-effectiveness and the ability to quantify microorganisms by counting Colony-Forming Units (CFUs) (Davis, 2014; Lopes et al., 2016).This technique also allows for the phenotypic analysis of colonies, facilitating the identification of the evaluated species. The Drop-plate technique, a variation of CFU counting, offers advantages such as reduced time and effort for analysis compared to the Spread-plate (Davis, 2014; Herigstad et al., 2001; Naghili et al., 2013). Conventional culture-dependent techniques also permit the use of selective and differential culture media, enabling the separate isolation and identification of different groups of yeasts and bacteria, such as an improved medium that was studied by Zúñiga and colleagues aiming at distinguishing between homofermentative and heterofermentative lactic acid bacteria (Zúñiga et al., 1993).

While culture-dependent methods like these are considered the gold standard, they present drawbacks, including long preparation times, difficulty in differentiating closely related species, extended incubation periods, and a lack of information on microbial species interactions in a complex community, focusing on the isolated growth of microorganisms (Vinderola et al., 2019). Since culture-dependent methods rely on microbial growth, quantifying only reproducing cells, it can lead to an underestimation of viable microorganisms, such as those with active metabolism but incapable of reproducing (Viable But Not Culturable - VBNC) (Vinderola et al., 2019).This ties into the complex concept of cellular viability: what defines a viable cell?(Kwolek-Mirek & Zadrag-Tecza, 2014).

Another limitation of culture-dependent methods is the “great plate count anomaly,” a term described and coined by Staley and Knopka in 1985, and first observed by Razumov in 1932 (Razumov, 1932), referring to the significant difference in the orders of magnitude between the number of cells from natural environments forming colonies in culture-based plating methods and the number of cells quantified by culture-independent methods (Staley & Konopka, 1985). Factors such as viability, cultivability, selective culture media, suboptimal cellular growth conditions (pH, temperature, oxygen availability), and ecological interactions (competition, mutualism, predation, etc.) contribute to only a small fraction of the microbial diversity in a sample being able to reproduce and form visible colonies (Connon & Giovannoni, 2002; Steen et al., 2019).

With the evolution of molecular biology and molecular techniques, methods like Polymerase Chain Reaction (PCR), immunology-based methods, and techniques such as bright-field and fluorescence microscopy, flow cytometry, and dielectric spectroscopy have become more prevalent for microorganism identification and quantification since the 1970’s (Davis, 2014; Defossa & Wagner, 2014; Deshmukh et al., 2016; Franco-Duarte et al., 2019; Lopes, Cristina, et al., 2016). These culture-independent methodologies offer advantages, such as the ability to quantify both cultivable and non-cultivable cells. While providing rich information about microbial community composition, some culture -independent methodologies, like sequencing-based methods, may struggle to distinguish between viable and non-viable cells.

Viability is frequently a crucial parameter monitored during bioprocess development, and traditional plating methods, despite dating back to the 19th century, remain valuable for viable cell enumeration (Kwolek-Mirek & Zadrag-Tecza, 2014). However, flow cytometry and molecular-genomic approaches are gaining traction as potential methods for bacterial viability measurement in probiotic products (Bagheripoor-Fallah et al., 2015; Wilkinson, 2018). Despite their advantages, these culture-independent methods can be expensive and sophisticated, limiting their routine use in academic research domains. It’s worth noting that when comparing cell levels using culture-dependent and independent techniques, differences in counts may arise because culture-independent methods don’t assess cellular viability. Comparative studies between enumeration methods for probiotic bacteria in food showed variations, emphasizing the significant impact of the chosen enumer ation method on the results (Lahtinen et al., 2006). Several studies explored the comparative analysis of total enumeration and viability of yeasts and bacteria individually, highlighting the pros, cons, and challenges of each method (Davis, 2014; Kwolek-Mirek & Zadrag-Tecza, 2014; Vinderola et al., 2019; Wang et al., 2021). However, many fermentation processes involve mixed cultures containing multiple microbial species, making it crucial to quantify both yeast and bacterial cells simultaneously. In the food industry, for instance, there are numerous products and processes where yeast and bacterial cells coexist, particularly lactic acid bacteria, exemplified by kefir (Garofalo et al., 2015).

In the context of Brazilian sugarcane biorefineries, the quantification of yeasts and bacteria is essential for monitoring both the cellular concentration of the main microorganism responsible for the fermentative process, the *Saccharomyces cerevisiae* yeast, and contaminating bacterial cells, mainly composed of lactic acid bacteria (Vieira & Fernandes, 2012). In these environments, the quantification of yeast and bacterial cells is performed multiple times a day. Consequently, methods requiring less training and time for obtaining results, as well as methods with lower associated costs for equipment, are commonly preferred (Beckner et al., 2011; Ceccato-Antonini,2018, 2021). On the other hand, in environments such as pharmaceutical industries, where the final product generally has higher added value and there is more investment in infrastructure, equipment, and operator training, methodologies with higher accuracy, specificity, and complexity are typically adopted, even though pure cultures are usually present. After we started the present work, a recent study demonstrated successful and consistent simultaneous enumeration of yeasts and bacterial cells in mixed cultures using a specific image-based method, emphasizing the need for further comparative studies across different quantitative methods for complex samples (Williamson et al., 2023).

The present work was initiated to perform a comparative assessment of currently employed techniques for quantifying yeast and bacterial cells, with a special focus on simultaneous quantification in mixed samples through a single analysis, without aiming to describe the different species present within the complex sample. The evaluation criteria included analysis and equipment costs, analysis time, detectable cellular diameter range, ease of operation, ease of result acquisition and analysis, potential related challenges, and, crucially, the capacity for simultaneous quantification of yeast and bacterial cells in the same analysis. The choice of equipment and techniques for this work was also based on their availability in our laboratory or in laboratories that agreed to collaborate on the research, considering the difficulty in finding partner laboratories that possess all the selected methodologies and equipment mainly due to their high value.

## MATERIALS AND METHODS

### Yeast strain and growth conditions

Dehydrated cells of *Saccharomyces cerevisiae* PE-2 (LNF Latino Americana, Rio Grande do Sul, Brazil) were reactivated in 50 mL Falcon tubes with 10 mL of YPD medium, previously sterilized at 121 °C for 20 minutes, at 200 rpm (Innova 4430 Shaker Incubator, New Brunswick), at 30 °C for 24 h. For yeast cultivation, liquid YPD medium composed of 10 g/L yeast extract (Merck Millipore, Germany), 20 g/L Dextrose (D-glucose) (Sigma-Aldrich, Missouri, USA), and 20 g/L peptone (Neogen, Michigan, USA) w as used, along with YPD-agar medium containing the same components, supplemented with 20 g/L agar (Kasvi, São José dos Pinhais, Brazil). After reactivation in YPD liquid, the industrial yeast strain was cultivated on YPD-agar and incubated in a bacteriological incubator (model 502, Fanem) at 30 °C for 48 h. After this period, within a laminar flow hood, cells from one of the colonies were scraped with a sterile inoculation loop and used to inoculate a sterile 50 mL Falcon tube containing 15 mL of YPD medium and incubated at 150 rpm and 30 °C (Innova 4430 Shaker Incubator).

### Bacterial strain and growth conditions

Dehydrated cells of the species *Lactiplantibacillus plantarum subsp. plantarum* was obtained from a local pharmacy (Ophicinales – R. Manoel Matheus, 890, Santa Rosa – Vinhedo, SP, Brazil). The cells were reactivated in 50 mL Falcon flasks with 10 mL of Man Rogosa Sharpe (MRS) medium, in static cultures in a bacteriological incubator (model 502, Fanem), at 37 °C for 24 h. For LAB cultivation, MRS medium (Neogen, Michigan, USA) was used (Peptone 10 g/L, yeast extract 5 g/L, meat extract 10 g/L, glucose 10 g/L, potassium phosphate 2 g/L, sodium acetate 5 g/L, magnesium sulfate 0.2 g/L, manganese sulfate 0.05 g/L, Tween 80 1.08 g/L, and ammonium citrate 2 g/L), along with MRS-agar medium containing the same components, supplemented with 20 g/L agar (Kasvi, São José dos Pinhais, Brazil). After reactivation in the MRS liquid medium, the strain *L. plantarum* was cultivated on MRS at 37 °C for 24 h (model 502, Fanem). After this period, within the laminar flow hood, cells from one of the colonies were scraped with a sterile inoculation loop and used to inoculate 50 mL Falcon flasks containing 15 mL of MRS medium and incubated at 37 °C statically.

### Drop-plate CFU counting method

The Drop-plate technique followed an adaptation of the protocol described by Herigstad et al. (2001). Yeast and LAB strains were individually cultivated as described before. Cells of *S. cerevisiae* and *L. plantarum* were separately cultivated in 50 mL Falcon flasks containing 10 mL of YPD and MRS media, at 30 and 37°C, respectively, for 24 h. From this pre-culture in a liquid medium, a cell suspension in a 1:1 proportion (yeast: bacteria) was created following a previous quantification of the yeast and bacterial cell suspensions through bright-field microscopy. Subsequently, serial dilutions were performed in 15 mL Falcon tubes containing 9 mL of 0.9% saline solution with one mL of each cell suspension (1:9 v/v dilution). This tube was vortexed (Vortex Mixer, K45-2820, Kasvi) for approximately 8 s, and 1 mL of its content was aliquoted and transferred to another tube subsequently. For plating, Petri dishes with ∼20 mL of YPD medium supplemented with chloramphenicol (100 μg/mL) (Inlab, São Paulo, Brazil) and MRS medium supplemented with cycloheximide (50 μg/mL) (Inlab, São Paulo, Brazil) were used, as a selective media for yeast and bacteria, respectively. Each Petri dish was divided into three parts, each reserved for one of the three dilutions selected. In each part, a total of 30 μL was dispensed in three separate drops (each drop containing 10 μL). Once the drops had dried on the agar medium, Petri dishes were inverted and incubated at 30 or 37°C (incubator model 502, Fanem) for 20 to 24 h. This procedure was conducted in triplicate.

### Spread-plate manual and automated CFU counting methods

Yeast and bacterial strains were cultivated and subsequently diluted as described in the Drop-plate methodology. For the plating step, Petri dishes with ∼20 mL of YPD-agar medium supplemented with chloramphenicol (100 μg/mL) and MRS-agar medium supplemented with cycloheximide (50 μg/mL) were used. One hundred microliters of the mixed cell suspension were added to the appropriate medium, spread using a Drigalski spatula, and incubated at 30 or 37°C for 24 to 48 h. The mixed cell suspension was evaluated at three dilutions and in quintuplicate. In addition to the manual counting of CFUs in the Spread-plate plating technique, an automatic count was also performed. For this purpose, each set of plates cultivated with the mixture of yeast and bacterial cells, as described previously, was placed in a chamber on a smooth black background at a fixed distance of 30 cm from the photographic device (Smartphone Samsung Galaxy A52), and images were captured in JPEG format for subsequent analysis. Based on the obtained images of each plate, a Python code was developed (Supplementary File 1) and used to count the colonies on each plate automatically.

### Direct counting through bright field microscopy method

Yeast and bacterial cells were quantified using an adaptation of a standard method used in some sugarcane biorefineries for counting rods, referred to as the “Slide method” as an alternative to the conventional hemocytometer method, such as the Neubauer Chamber, due to the challenges of quantifying bacterial cells with such approaches (Ceccato-Antonini, 2021). This adaptation involved the addition of centrifugation steps for the cell suspension and resuspension in a sterile 0.9% saline solution to remove the culture medium. Additionally, the Nile blue/Methylene blue viability stain was replaced by the Erythrosine B stain.

For cell counting using the adapted methodology, aliquots of cell suspensions from the *S. cerevisiae* PE-2 and *L. plantarum* strains, previously cultured, were taken from their respective 50 mL Falcon tubes and stored in 2 mL Eppendorf tubes. The tubes were centrifuged (MiniSpin, Eppendorf – Hamburg, Germany) at 4800 g for 10 min at room temperature. The supernatant was removed, and the cell pellet was resuspended in 1 mL of sterile 0.9% saline solution. Another centrifugation step was performed (under the same conditions), where the supernatant was removed, and 1 mL of sterile 0.9% saline solution was added to remove the culture medium from the cell suspensions as much as possible. For the simultaneous counting of yeast and bacterial cells, a cell suspension of yeast and bacteria in an approximate 1:1 rat io was prepared. The cell counting of yeast and bacteria was performed using the Eclipse E200 microscope (Nikon, Tokyo, Japan) with an attached Pri meCam camera (Prime HD model, Florida, United States), using a glass slide measuring 26.0 x 76.0 mm (thickness ∼1 to 1.2 mm) and a cover glass measuring 20 x 20 mm. A 400x objective lens was used for yeast counting, while a 1000x objective lens with immersion oil was used for bacteria quantification. Before adding the samples of the cell mixture for microscope counting, they were diluted with a 0.9% (w/v) NaCl solution to achieve a concentration of approximately 10^7^,cells/mL for each cell population (first yeast cells were counted, and then bacterial cells). The cells were manually counted in quintuplicate. The determination of cell concentration was given by Equation 1, described in the Slide Method. (Ceccato-Antonini,2012)

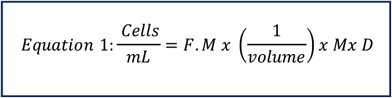

Where: M = total viable rods ÷ Number of counted fields; D = dilution used; F.M. = Microscope Factor.

The microscope factor (F.M.) corresponds to the ratio between the area of the coverslip used in the analysis and the area of the field of view of the objective lens used in the microscope. In other words, it is the number of visual fields that fit in the area underneath the cover slip. The microscope field of view area is determined by the software after calibration of the camera attached to the microscope with the aid of a glass slide with a microscope calibration ruler.

For determining cell viability using the microscopy technique, the Erythrosine B cell viability stain (0.1%) was added to the cell mixture suspension in PBS buffer, pH 7.2 (Dihydrogen phosphate 26.22 g/L, 7.78 g/L Sodium carbonate, Magnesium chloride 5 mM). This step was done in a 1:1 v/v ratio. Cell suspensions were incubated with the stain for 5 min previous cell enumeration. Cells stained with Erythrosine B acquired a pink color and were considered non-viable. Cell viability was determined by the ratio of the number of viable cells to the total number of cells in each sample.

### Flow cytometry method

Yeast and bacterial cells were individually cultivated as described previously, and samples were aliquoted into 2 mL Eppendorf tubes with 1.5 mL of cell suspension. These samples were then filtered through a 50 μm filter (CellTrics, Sysmex, Kobe, Japan), vortexed (Vortex Mixer, K45-2820, Kasvi) for 10 s, and analyzed on the Attune NxT flow cytometer, Thermofisher (Waltham, USA). Initially, only cell concentration was evaluated without considering cell viability. For the determination of the absolute number of microbial cells in mixed samples, cell suspensions from isolated yeast and bacterial cultures were analyzed and their cell concentration was determined. From this previous step, once the cell concentration of each yeast and bacterial cell population was known, the mixed cultures in the proportions of 1:1 and 100:1 of yeast and bacterial cells were prepared, and the simultaneous enumeration of yeast and bacterial cells was determined. The quantification of yeast and bacterial cells was performed by differentiating their sizes on the flow cytometer, in collaboration with Dr. Ana Paula Jacobus from Unesp Rio Claro, at the Institute of Bioenergy Research (IPBEN). The quantification of the yeast and bacterial cell mixture was performed in quintuplicate.

For determining cell viability using flow cytometry, cell suspensions were incubated with propidium iodide (PI) solution at 5 .7 μg/mL (18 μL of a 100 μg/mL PI stock solution in 300 μL of cell suspension) for 1 minute in the absence of light and then analyzed on the same above-mentioned flow cytometer.

### Coulter Counter method

The Multisizer IV Coulter Counter equipment (Beckman, Brea, California, United States) provides number, volume, mass, and size distribution measurements in a single analysis, with a particle diameter range from 0.4 μm to 1,200 μm. For determining the cell concentration of co-culture samples using this equipment, the volumetric mode was selected. Once the volume of the sample to be analyzed was input, the absolute value of quantified particles was provided, thereby obtaining the cell concentration. In summary, yeast and bacterial cells were cultivated individually, and their samples were aliquoted into 2 mL Eppendorf tubes with 1.5 mL of cell suspension, vortexed, and analyzed. In this analysis, for the determination of the absolute number of microbial cells, samples from isolated yeast and bacterial cultures were analyzed, followed by the mixed culture. The simultaneous quantification of yeast and bacterial cells was performed in quintuplicate, using a glass tube with an opening aperture of 20 μm.

### Systematic Study

For the comparative study of the selected techniques for the simultaneous quantification of yeast and bacterial cells, the strains of *S. cerevisiae* PE-2 and *L. plantarum* were individually cultivated as described before and 1 mL aliquots from each culture were withdrawn and stored in 1.5 mL Eppendorf tubes. In total, 8 aliquots of 1 mL each were taken from each culture, with 2 mL allocated for each technique to be used. For the flow cytometry and Multisizer IV automatic counter techniques, 1 mL of the cell suspension of each microorganism was used for the initial separate quantification of yeast or bacteria, while the other 1 mL was used to prepare the mixed cell suspension. On the other hand, for the bright-field microscopy and plating techniques, Spread-plate and Drop-plate, only 1 ml of the cell suspension of each microorganism was used for the preparation of the mixed culture and subsequent simultaneous quantification steps. As the use of the flow cytometer and Multisizer IV was done in collaboration with different facilities, the analyses were not conducted on the same day but according to the following schedule (Table 1).

**Table 1.** Schedule of activities for experiments related to the systematic comparative study of techniques for the simultaneous quantification of yeast and bacterial cells.

| Day | 1<sup>st</sup> day- Monday | 2<sup>nd</sup> day - Tuesday | 3<sup>rd</sup> day- Wednesday | 4<sup>th</sup> day- Thursday | 5<sup>th</sup> day- Friday |
| --- | --- | --- | --- | --- | --- |
| Activities | i) *S. cerevisiae* (PE-2) cultivation in YPD media at 30 °C for 24h;<br>ii) *L. plantarum* cultivation in MRS media for 37 °C for 24h | i) Plating techniques (Drop-plate and Spread-plate);<br>ii) Cell enumeration by bright-field microscopy | i) Manual CFU counting (Drop-plate method);<br>ii) Cell enumeration by Flow Cytometry | Cell enumeration by Coulter Counter Principle (Multisizer IV) | Manual and automated CFU counting (Spread-plate method) |
| Location | Campinas- SP, Brazil | Campinas- SP, Brazil | Rio Claro- SP, Brazil | São Bernardo -SP, Brazil | Campinas- SP, Brazil |

### Data Analysis

#### Statistical analysis

The means and (sample) standard deviations of yeast and bacterial cell counts for each evaluated technique were calculated using Microsoft Excel 2023 software (Redmond, USA), considering the quintuplicates of each assay.

The statistical analysis of the comparative study between the evaluated techniques in this work was performed using Origin Pro 8.5 software. An Analysis of Variance (ANOVA) was conducted, and Tukey’s test was applied pairwise for the techniques.

#### General Comparison Between Cell Quantification Techniques

Regarding the comparative study between cell quantification techniques, a table was elaborated with the following information: analysis and equipment costs, analysis time, detectable cellular diameter range, ease of operation, the possibility of storing samples for later analysis, and, most importantly, the ability to quantify yeast and bacteria simultaneously.

## RESULTS

In this work, cell suspensions were prepared from cultivating yeast and bacterial cells in YPD and MRS media, respectively. F rom these suspensions, aliquots were withdrawn and directly used or stored for quantification using different techniques, as outlined in the experimental plan presented in Table 1. For plating techniques (Spread-plate and Drop-plate) and bright-field microscopy, mixed suspensions of *S. cerevisiae* yeast and *L. plantarum* bacteria populations were prepared in a 1:1 cell concentration ratio and analyzed. This ratio corresponds to what is observed during acute contamination episodes in Brazilian sugarcane biorefineries, in which both yeast and bacterial cells might reach concentrations in the order of 10^8^,cells/mL (Ceccato-Antonini, 2021). As for flow cytometry and Coulter Counter techniques, using the Multisizer IV equipment, mixed suspensions were prepared in ratios of 1:1 and 100:1 and analyzed. Additionally, the yeast and bacterial cell populations were quantified separately using all these techniques. A proportion of 100:1 yeast-to-bacterial cells is typical in such biorefineries under normal processing conditions (or chronic contamination).

### Validation of the Slide Method

Yeast and bacterial cells were quantified using an adaptation of a standard method used in some sugarcane biorefineries for counting rods, referred to as the “Slide method”. An initial validation step was performed by comparing the enumeration of yeast cells through the “Slide Method” and a Neubauer Chamber method (Table S1 and Fig. S1).

### The conventional techniques, plating and microscopy, showed higher data variability

For yeast cell enumeration from a mixed sample, it is observed that the Drop-plate technique and microscopy presented the highest average values of yeast cell concentration (Table 2). The cell concentrations obtained through these techniques were nearly double those obtained by Spread-plate plating techniques, whether through manual or automatic counting, which in turn yielded values very close to each other. Regarding the standard deviation of each technique, in terms of the coefficient of variation, it can be observed that the Spread-plate technique exhibited the highest variation among its replicates compared to the other techniques. From the data in Table 2 (coefficients of variation), it is noted that there is less variation among the results obtained by different techniques for determining bacterial cell concentration than for yeast cells, with the exception of the microscopy technique. When comparing the techniques evaluated in Table 2, the microscopy technique for bacterial cell enumeration led to the highest CV, being the least precise technique among those evaluated in this study up to this point. It is also worth noting that for both yeast and bacterial cell quantification, the Drop-plate technique presented lower CV values when compared to the Spread-plate techniques.

**Table 2.** Cell concentration values obtained for yeast and bacterial cells with plating techniques (Drop-plate and Spread-plate) and microscopy*.

|  | Technique | Unit | Rep1 | Rep2 | Rep3 | Rep4 | Rep5 | Mean | s.d | Coefficient of variation (%) |
| --- | --- | --- | --- | --- | --- | --- | --- | --- | --- | --- |
| *S. cerevisiae* (PE-2) | Drop-plate | CFU/mL x10<sup>-7</sup> | 9.0 | 7.0 | 7.5 | 8.5 | 7.0 | **7.80** | 0.91 | 11.7 |
|  | Spread-plate | CFU/mL x10<sup>-7</sup> | 5.2 | 3.4 | 4.0 | 3.2 | 5.0 | **4.16** | 0.91 | 21.9 |
|  | Automated Spread-plate | CFU/mL x10<sup>-7</sup> | 4.8 | 5.2 | 3.6 | 3.4 | 4.0 | **4.20** | 0.77 | 18.3 |
|  | Microscopy | cel/mL x10<sup>-7</sup> | 9.4 | 8.3 | 7.5 | 9.6 | 6.12 | **8.18** | 1.43 | 17.5 |
| *L. plantarum* | Drop-plate | CFU/mL x10<sup>-9</sup> | 2.30 | 2.20 | 2.30 | 2.05 | 2.00 | **2.17** | 0.14 | 6.4 |
|  | Spread-plate | CFU/mL x10<sup>-9</sup> | 2.36 | 2.28 | 2.56 | 2.08 | 1.68 | **2.19** | 0.33 | 15.1 |
|  | Automated Spread-plate | CFU/mL x10<sup>-9</sup> | 2.16 | 2.00 | 2.32 | 1.92 | 1.62 | **2.00** | 0.26 | 13.0 |
|  | Microscopy | cel/mL x10<sup>-9</sup> | 2.53 | 3.22 | 2.24 | 2.71 | 1.32 | **2.40** | 0.70 | 29.2 |

\*Yeast and bacterial cell suspension in a 1:1 ratio

In addition to the manual counting of Colony-Forming Units (CFUs) of yeast and bacteria in the Spread-plate technique, automatic counting was also performed using a computer code developed for this purpose, as described in the Materials and Methods section and Supplementary Material (Supplementary File 1 and Fig. S2).

### Flow Cytometry is the most precise technique for the simultaneous enumeration of yeast and bacteria at 1:1 ratio

In the flow cytometry technique, the mixed suspension of yeast and bacteria was quantified in the ratios of 1:1 and 100:1 (Fi g. 1), aiming to represent conditions of acute and standard contamination in industrial fermentation vessels, in sugarcane biorefineries. The order of magnitude for yeast concentrations is 10^8^,cells/mL, while for contaminating bacteria, the order of magnitude is 10^6^,cells/mL under normal process conditions, rising to up to 10^7^,or even 10^8^,cells/mL during episodes of acute contamination. (Ceccato-Antonini, 2021) The yeast cell concentrations, obtained in quintuplicate, are presented in Table 3 for each studied cell ratio condition.

**Table 3.** Cell concentration values of yeast and bacterial suspensions obtained from the flow cytometry technique at yeast:bacteria ratio of 1:1 and 100:1.

|  |  | Cell concentration |  |  |  |  |  |  |  |
| --- | --- | --- | --- | --- | --- | --- | --- | --- | --- |
|  | Technique | Rep1 | Rep2 | Rep3 | Rep4 | Rep5 | Mean | s.d | CV (%) |
| *S. cerevisiae*<br>($\text{cel/mL} \times 10^{-7}$) | Flow cytometry (Ratio 1:1) | 3.28 | 3.21 | 3.37 | 3.52 | 3.63 | **3.40** | 0.17 | 5.0 |
|  | Flow cytometry (Ratio 100:1) | 3.34 | 2.90 | 2.85 | 2.86 | 2.99 | **2.99** | 0.20 | 6.7 |
| *L. plantarum*<br>($\text{cel/mL} \times 10^{-9}$) | Flow cytometry (Ratio 1:1) | 1.50 | 1.35 | 1.39 | 1.71 | 1.42 | **1.47** | 0.14 | 9.5 |
|  | Flow cytometry (Ratio 100:1) | 7.06 | 2.22 | 1.64 | 2.44 | 2.42 | **3.16** | 2.21 | 69.9 |

Fig. 1a shows the yeast population within the region defined as gate R2 in the histogram graph (Count x FSC) and R 3 in the dot-plot graph (SSC x FSC), while the bacterial population is within the region defined as gate R1 in the histogram graph (Count x FSC) and R4 in the dot-plot graph (SSC x FSC) in Fig. 1b. Fig. 1c represents the analysis of the suspension containing the mixture of yeast and bacteria in a 1:1 ratio, while Fig. 1d represents the suspension containing the mixture of yeast and bacteria in a 100:1 ratio. The yeast cells of the mixture are in gates R2 and R3, while the bacterial cells are in gates R1 and R4. As observed in Fig.1a and b, yeast cell debris occupies the same region as bacterial cells, highlighted by the red rectangle. This may lead to signal overlap between yeast cell debris and bacterial cells, resulting in an overestimation of bacterial cell quantification. In both cases, the quantification of yeast cells in a mixed sample by the flow cytometry technique showed standard deviations representing low coefficients of variation, between 5 and 7%, in a 1:1 and 100:1 ratio respectively. It is noteworthy, from the flow cytometry results, that while bacterial cell quantification in a mixed sample is possible using this technique when the yeast-to-bacterial cell ratio is around 1:1, the same does not hold when the ratio is 100:1. This is evident from the high standard deviation (coefficient of variation of approximately 70%) presented in Table 3. These results highlight the difficulty of differentiating between bacterial and yeast cell debris by the flow cytometry technique at a 100:1 ratio, due to their similar sizes, and raise the question of whether there is an alternative methodology capable of making this differentiation using this or other techniques.

**Fig. 1.**
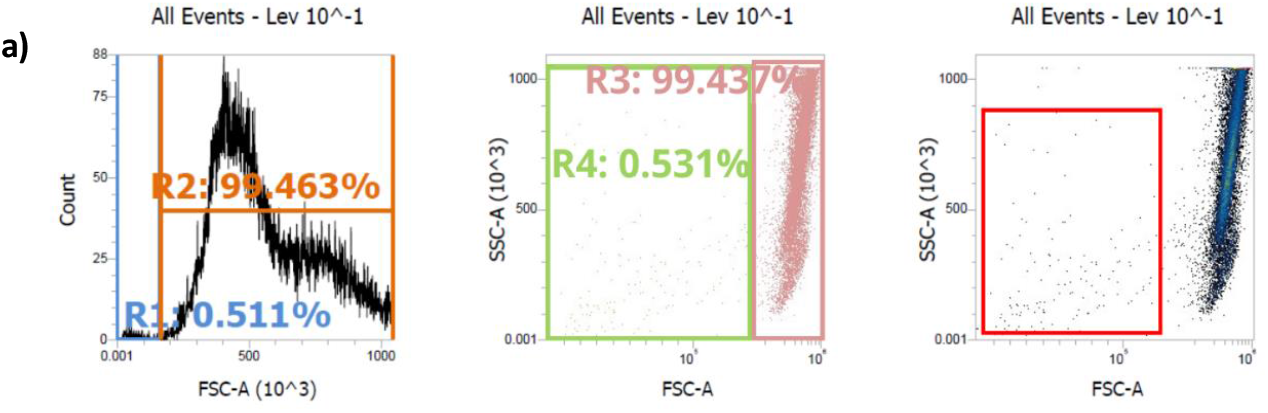
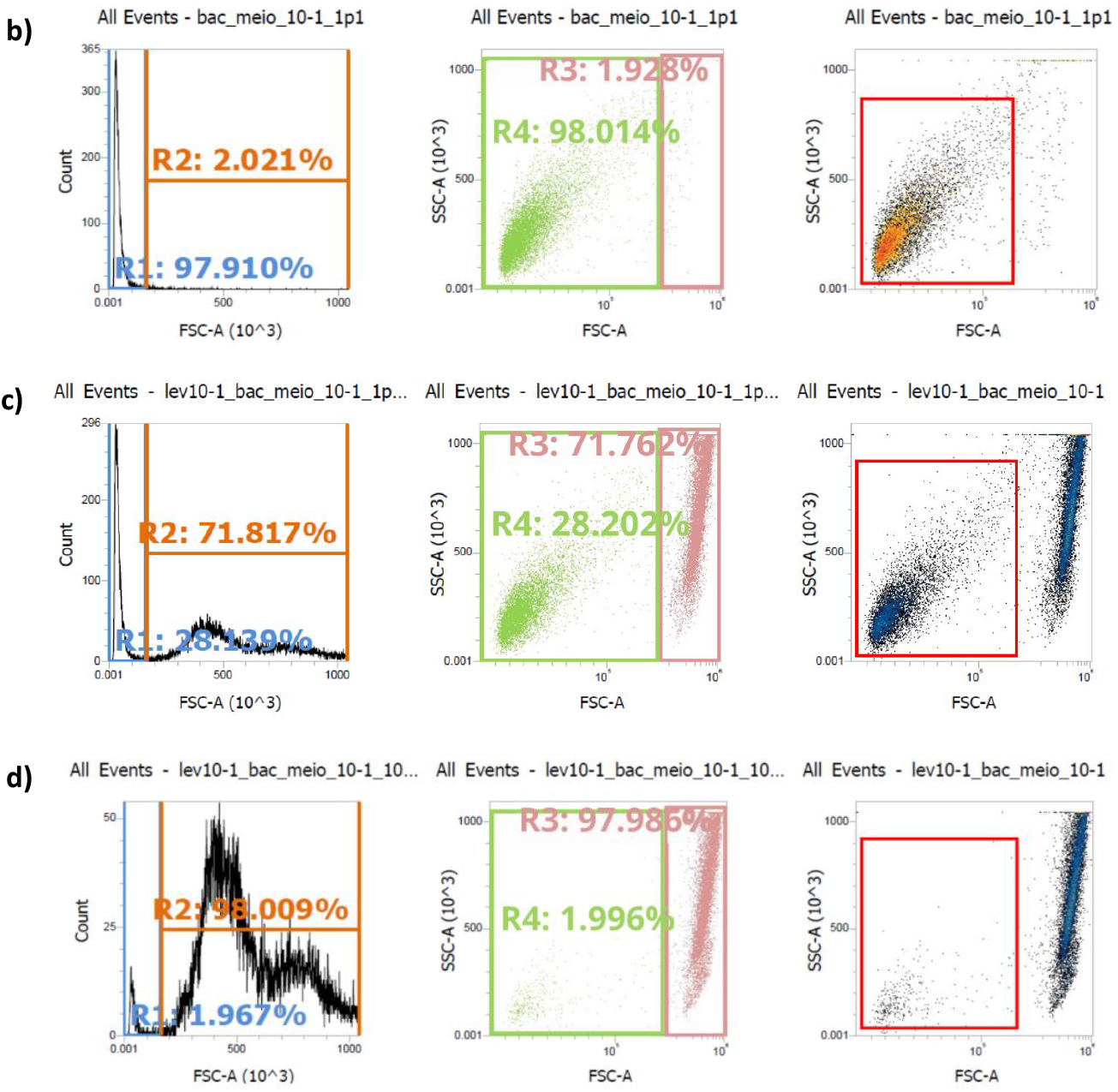
Cell quantification results by flow cytometry a. Histogram and Dot Plot of cultures prepared in suitable culture media for a) *S. cerevisiae*; b) *L. plantarum*; c) mixed culture of *S. cerevisiae* and *L. plantarum* in a 1:1 ratio; d) mixed culture of *S. cerevisiae* and *L. plantarum* in a 100:1 ratio; Gates R1 and R4 represent the region of the *L. plantarum* population; and gates R2 and R3 represent the region of the *S. cerevisiae population*.

### The Coulter counter technique not only allows simultaneous enumeration of yeasts and bacteria but also cell diameter range determination

The mixed suspension of yeast and bacteria was also quantified using the Coulter Principle with the Multisizer IV equipment. The results obtained for the simultaneous quantification of yeast and bacterial cell populations are presented in Fig. 2.

**Fig. 2.**
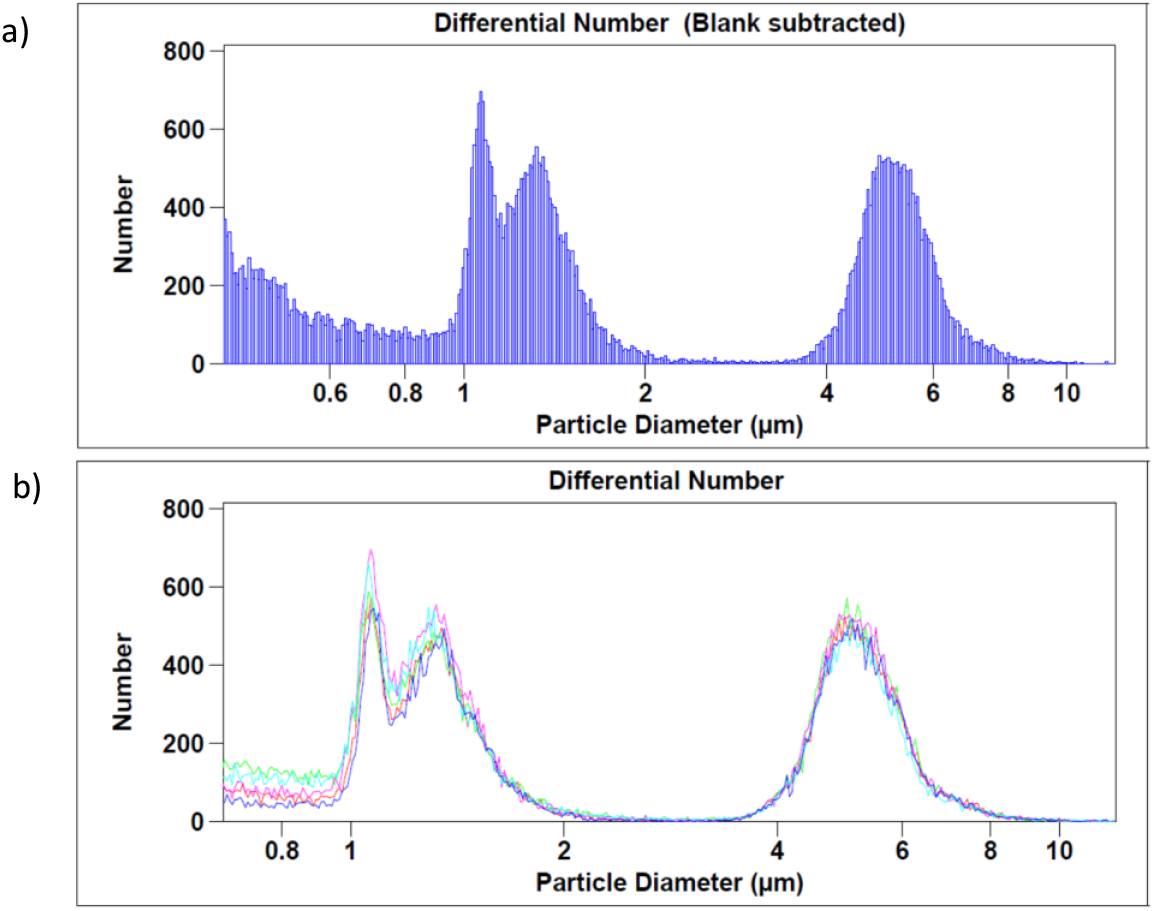
a) Histogram of populations of *S. cerevisiae* and *L. plantarum* prepared from cultivation in YPD and MRS media, respectively, and subsequent mixing (in a ratio of approximately 1:1) followed by analysis on the Multisizer IV. b) Compilation of the quintuplicate histograms obtained for the cell suspension of *S. cerevisiae* and *L. plantarum* on the Multisizer IV.

From the data presented in Fig. 2, it is noted that the yeast population has an average cell diameter between 5 and 5.5 μm, approximately, which corroborates with data from the literature (Fukuda, 2023). For yeast quantification with this technique, the interval of 3 to 11 μm was arbitrarily selected. The bacterial population, on the other hand, exhibits two overlapping subpopulations with an average cell diameter of around 1 and 1.8 μm. For the subsequent systematic study carried out in this work, the interval of 0.85 to 2.2 μm was arbitrarily defined for bacterial cell quantification.

The quantification of both yeast and bacterial cells from a mixed suspension could be performed on the Multisizer IV equipment at a 1:1 ratio, with a coefficient of variation equivalent to approximately 5% and 9%, respectively. However, it was not possible to distinguish between bacterial cells and yeast cell-derived debris when the suspension was in the order of 100:1.

### Yeast and Bacterial Cells in Mixed Samples can be Simultaneously Enumerated by Different Techniques

This work aimed to conduct a comparative analysis of the selected techniques for the simultaneous quantification of yeast and bacterial cells in the same sample. For statistical analysis of the results, an Analysis of Variance (ANOVA) was performed using Origin Pro 8.5 software, yielding a p-value of 5.4×10^-12^,for the comparative analysis of all techniques in yeast cell quantification and 7.1×10^-5^,for the comparative analysis of all techniques in bacterial cell quantification. These p-values indicated that there is at least one mean that differs significantly from the others in the evaluated groups. Subsequently, a Tukey’s post hoc test was performed to identify which groups have significantly different means after detecting these significant global differences by ANOVA. Tukey’s post hoc test makes pairwise comparisons, and the p-values obtained are presented in the supplementary material (Tables S2 and S3).

The p-values obtained for pairwise analyses of each technique in this study show that, for the quantification of yeast cells in a mixed sample with bacterial cells, the Drop-plate plating and bright-field microscopy techniques are statistically equivalent to each other but different from the other evaluated techniques. The Spread-plate with manual and automatic counting, flow cytometry in both 1:1 and 100:1 ratios, and the Coulter Counter Multisizer IV are statistically equivalent to each other for the yeast cell enumeration. For the quantification of bacterial cells in a mixed sample with yeast cells, the p-values obtained for pairwise analyses of each technique show that the Coulter Counter technique with the Multisizer IV equipment is statistically different from the other evaluated techniques. When compared to flow cytometry at the 100:1 ratio (which showed a coefficient of variation of 70%), the Coulter Counter technique had a p-value of 0.07634. This means that, depending on the chosen significance level, these techniques may still be considered statistically equivalent or not (e.g., with a significance level of 0.1). On the other hand, Drop-plate technique, Spread-plate with manual and automatic counting, bright-field microscopy, and flow cytometry in both 1:1 and 100:1 ratios (despite having a high coefficient of variation) are statistically equivalent to each other. The comparison between flow cytometry at the 1:1 and 100:1 ratios yielded a p-value of 0.0853, and the same interpretation as for the Multisizer IV and flow cytometry at 100:1 can be applied to this analysis.

The results of yeast and bacterial cell concentrations from a mixed culture obtained in quintuplicate for each evaluated technique in this study, including Drop-plate, Spread-plate with manual counting, Spread-plate with automatic counting, microscopy using the Slide Method adaptation, flow cytometry (at yeast-to-bacterial cell ratios of 1:1 and 100:1, respectively), and automatic counting using the Coulter Principle (Multisizer IV), were grouped in Fig. 3.

**Fig. 3.**
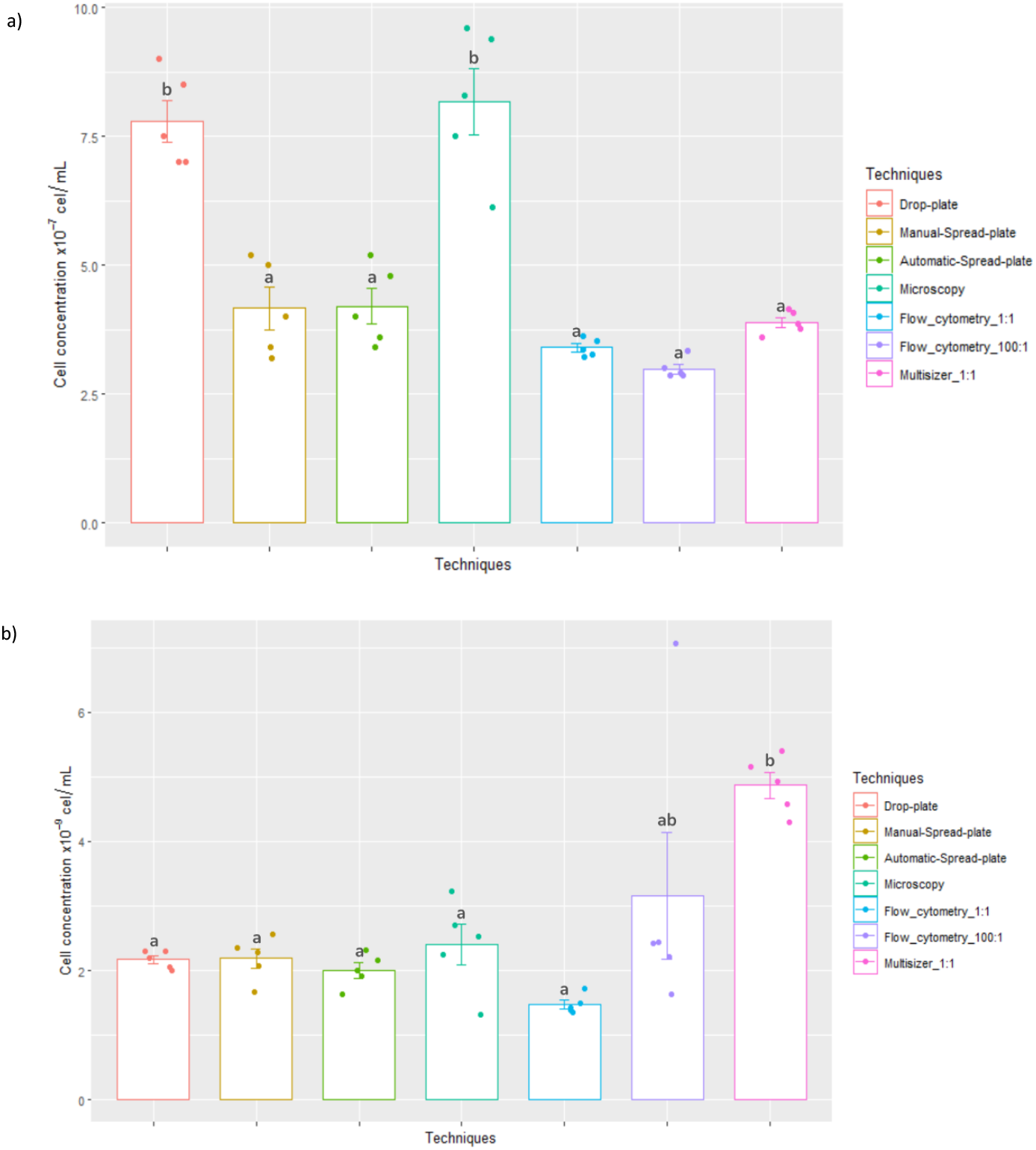
a) Yeast cellular concentrations obtained from mixed culture (with *L. plantarum* bacteria) using different cell quantification techniques; b) Bacterial cellular concentrations obtained from mixed culture (with *S. cerevisiae* yeast) using different cell quantification techniques. Error bars represent standard deviations from analyses conducted in quintuplicate.

Finally, comparing the determination of yeast and bacterial cell viability by bright-field microscopy and flow cytometry in this study, the determination of cell viability for yeast and bacterial cells by bright-field microscopy using the Erythrosine B viability dye was 91.6% and 88.80%, respectively, with coefficients of variation of 2.9% and 3.1% (Table S4). In the flow cytometry technique, the yeast cell viability was 90.4% with a coefficient of variation of 0.9%, while for bacterial cells, it was 99% with a coefficient of variation of 0.7% (Table S5).

## Discussion

In this work, a comparative analysis was conducted between different techniques for the simultaneous quantification of yeast and bacterial cells in the same sample. Simultaneous quantification of yeast and bacterial cells poses several challenges due to the difference in size, morphology, and biochemical characteristics of these microorganisms. Yeast cells are typically larger (5 to 15 μm in diameter, depending on genus and species, with oval/spherical shapes) and have more complex morphology and physiology, including cellular organelles, compared to bacterial cells (Horst Feldmann, 2012a). Bacterial cells, on the other hand, are usually smaller and exhibit various shapes (including cocci, bacilli, and spirals), with their size also varying by genus, making the variability in size and shape challenging for the development and/or use of a single quantification method that works well for both yeast and bacterial cells (Michael T. Madigan, 2016). Additionally, for culture-dependent methods, yeast, and bacterial cells may require different culture conditions (e.g., growth medium, temperature, pH, among others), further complicating the scenario for cultivating both microorganisms in the same environment, since incompatibilities in culture conditions can impact cell viability and growth rates (Horst Feldmann, 2012b). Moreover, there is the issue of interference signals and overlap between these microorganisms. Yeast and bacterial cells can interfere with each other in some quantification methods, such as flow cytometry, where the presence of one cell type can lead to overlapping signals, making the precise distinction of these two populations challenging, as seen in the overlap of signals from yeast cellular debris with signals from bacterial cells by flow cytometry technique. In the bright-field microscopy methodologies, along with automated counters that use the microscopy principle, the challenge of simultaneously quantifying yeast and bacterial cells is due to not only their difference in size but also the limitation of the method itself in enabling sufficient resolution to identify and quantify small cells and particles, such as the bacterial cells. Finally, there is the issue of standardizing the chosen method, as establishing a method to quantify yeast and bacterial cells simultaneously in the same sample can be a considerable challenge.

The comparative analysis of the results obtained by yeast cell quantification methods in a mixed sample, in the present work, gathered in Fig. 3a and Tables 2,3,4 and S2, show that when statistically analyzed in pairs, the results of the techniques had p-values greater than 0.21427, except for the Drop-plate plating and microscopy techniques. When these two techniques were compared individually with the others, the p-values were lower than 0.05. This result indicates that, in the present study and under the conducted experimental conditions, the results from these two latter techniques (with a p-value of 0.99 between them) statistically differed from the others. Regarding the reproducibility of the evaluated techniques in this study, it can be observed from the analysis of the coefficient of variation values of the replicates that flow cytometry and Coulter Counter, with the Multisizer IV equipment, are the ones that presented the most reproducible results when compared to plating and microscopy techniques in the quantification of yeast cells. Additionally, when comparing the results obtained by yeast cell quantification using the Spread-plate plating technique with manual and automatic counting, it was possible to obtain a p-value equivalent to 1 calculated between these two methodological options, indicating that these methodologies can be considered statistically equivalent. Choosing automated image analysis for counting allows a reduction in the analysis time, at least concerning colony counts. Moreover, it reduces the analyst’s interference in the results.

**Table 4.**
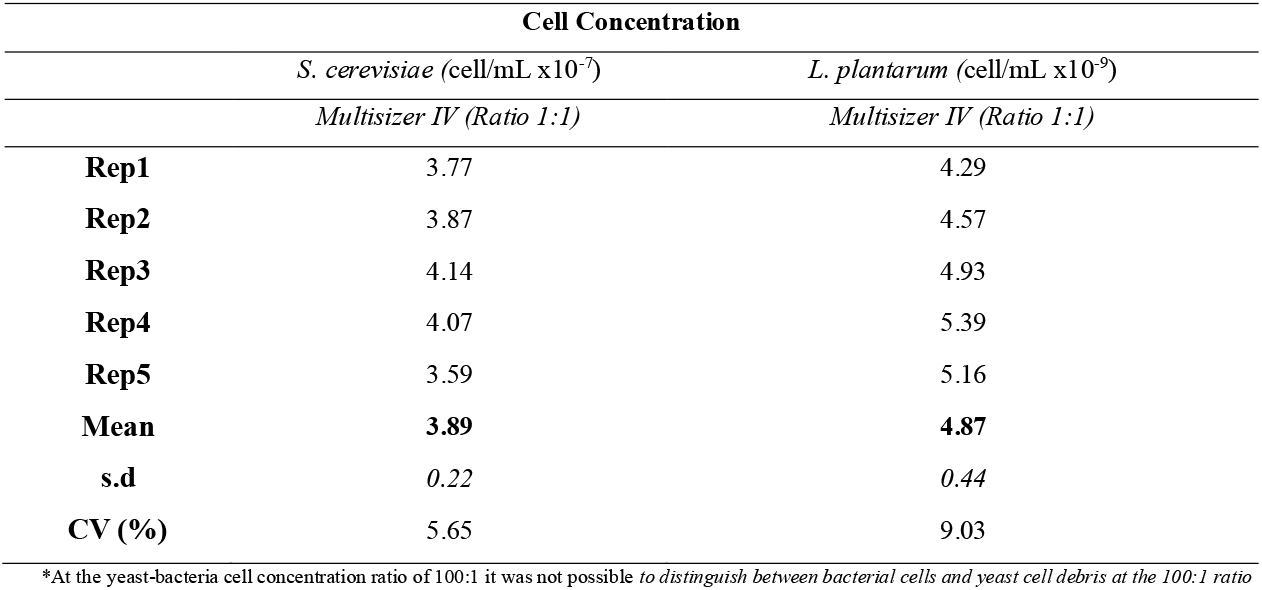
Yeast and bacterial cell concentration values obtained by Multisizer IV at yeast to bacteria ratios of 1:1.

On the other hand, the comparative analysis of the results obtained by bacterial cell quantification methods in a mixed sample, gathered in Fig. 3b and Tables 2,3,4 and S3, show that when analyzed in pairs, the techniques had p-values greater than 0.67, except for the Coulter Counter techniques with the Multisizer IV equipment and flow cytometry at a 100:1 ratio of yeast and bacteria. For the bacterial cell quantification from the mixed culture, it was observed that the Multisizer IV presented approximately twice the average value of the cell concentration compared to the other techniques. One possible explanation for this result is that part of this increase in bacterial cell concentration may have originated from the overlap of signals from bacterial cells with yeast cell debris, which occupy the same region of cell diameter (between 0. 85 and 2.2 μm). Additionally, in this technique, it was observed that the average size of the bacterial population was higher than observed previously during the preliminary testing stages of the methodologies, where cell suspensions were prepared by direct hydration of dried *L. plantarum* cells (data not shown). Besides the increase in average size (from 0.75 to 1.8 μm to 0.85 to 2.2 μm), the emergence of two distinct but close-sized populations of bacterial cells was also observed (Fig. 2). This possibly occurred due to the liquid culture step performed before the analyses in the different techniques evaluated here. Additionally, the *L. plantarum* species can form short “chains” of adjacent cells, as highlighted in Supplementary Fig. S3, which may have led to the average increase in cell diameter and the two populations observed in Fig. 2.

Regarding the flow cytometry technique, the quantification of bacteria from a mixed sample proved to be efficient with a low standard deviation at a 1:1 ratio with yeast cells. However, at a 100:1 ratio, a much higher standard deviation was observed (coefficient of variation of 69.9%) among replicates in this technique, possibly also caused by the overlap of yeast cell debris with bacterial cells. It is important to highlight the impact of an adequate gate selection in this technique, which emphasizes a trained operator since this step is done manually for each analysis. In the bright-field microscopy technique, as observed for the quantification of yeast cells, a higher standard deviation (with a coefficient of variation of 29%) in bacterial quantification was noticed. Although it presented a high coefficient of variation, the result obtained for bacterial quantification by bright-field microscopy supports the reported difficulty in quantifying this microorganism by the bright-field microscopy method, even though this methodology underwent some adaptation in this study. The bright-field microscopy technique, although widely used in different contexts for quantifying yeast and bacterial cells, mostly from pure cultures, has several limitations and characteristics leading this technique not only to require adequate training but also to be highly dependent on the analyst. Due to these factors, it may present errors of up to 30% associated with its protocol. These errors are often associated with operator errors, such as interpretation, especially regarding the viability stain’s color tones.

In their published work in 2015, Thomson et al. conducted a comparative study between traditional methods and a new automatic counting method with the Countstar equipment (Aber Countstar Yeast model - Aber, Aberystwyth, Wales) for yeast cell quantification and viability determination (Thomson et al., 2015). In their work, in addition to comparing yeast cell quantification by bright-field microscopy with the use of a hemocytometer and an automatic counter, Thomson and colleagues studied the impact caused by changing the operator during the counting by these two methodologies. Firstly, it was observed that the bright-field microscopy technique with a hemocytometer presented a higher standard deviation than that obtained by the automatic counter, with a coefficient of variation between 17 and 20% compared to that obtained by the automatic counter, varying between 4 and 6%. This observed difference, according to the authors, was expected since the Countstar automatic counter automates the highly error-prone manual counting procedure, giving rise to the known inter-operator differences in measurement by this methodology. In the same technique, in the same study, the determination of cell viability presented a coefficient of variation equivalent to 1.6%. Regarding the evaluation of the impact of inter-operator analyses, the authors observed that for obtaining cell viability, the change of the operator has a greater impact and generates more significant differences in the hemocytometer method than in the automatic quantification method. Furthermore, the cell viability parameter, in the bright-field microscopy technique, is a factor that, in addition to the errors associated with inter-sample and inter-operator variation, contributes to the subjectivity of this technique and the increase in variations between replicate analyses. This occurs because, regardless of the viability stain chosen, the process of stain internalization by non-viable/compromised cells may present different nuances in their coloration, depending on the physiological state of the cell and the mechanism of action of the chosen viability stain. The concept of cell viability and vitality itself, in this context, contributes to making this bright-field microscopy technique for determining cell viability dependent not only on the protocol and the stain used but also on the analyst for decision-making regarding the different tones that stained cells may obtain.(Kwolek-Mirek & Zadrag-Tecza, 2014)

Concerning the plating techniques, although starting from a single mixed cell suspension of yeast and bacteria, this technique cannot be considered a possibility for the simultaneous quantification of these microorganisms. This is because it is necessary to prepare and use different culture media, selective media (YPD with antibiotic and MRS with antifungal), so that the different populations studied in this work, the yeast *S. cerevisiae*, and the lactic acid bacteria *L. plantarum*, can grow separately. This fact constitutes a limitation of this technique, as it requires more time to obtain results (around 24 to 72 h, depending on the species/strain evaluated), is more labor-intensive, and provides the quantification only of viable and cultivable cells, those capable of growing, which can mask the absolute cell concentration of a microbial culture depending on the physiological conditions of the cells in question.

On the other hand, flow cytometry and Coulter Counter techniques, although showing some limitations such as the overlap of signals from bacterial cells with yeast cell debris, as they share the same size range, are capable of simultaneously quantifying these microorganisms in a single analysis. The reproducibility of these techniques is another positive point to be highlighted, along with ease of operation and result acquisition, despite the analysis of results being more complex and requiring training, especially for the flow cytometry technique. Additionally, flow cytometry allows obtaining the cell viability parameter, along with numerous other parameters if desired, making it a very versatile technique. Although it does not allow the determination of cell viability, the Coulter Counter technique allows obtaining not only cell concentration but also the size distribution of a population or particle, increasing the versatility of this equipment. However, both techniques require a filtration step for the cell suspension, especially if the 20 μm orifice tube is used and if less “clean” samples are used. Therefore, there is to some extent a limitation to their use or, at the very least, the development of appropriate protocols for different samples.

More recently, Williamson and colleagues (2023) developed a method based on image for the simultaneous quantification of *L. plantarum* and *S. cerevisiae* in a mixed fermentation culture. In their study, the authors used the image cytometry method with the Nexcelom Cellometer X2 equipment (Lawrence, MA, USA) (Williamson et al., 2023). They conducted three experiments: i) quantification of cell concentration in yeast and bacterial monocultures; ii) quantification of cell concentration in the mixed culture at various proportions; iii) monitoring of a mixed Berliner fermentation culture (a symbiotic fermentation process used to create Berliner Weisse beer, employing both yeast and lactic acid bacteria). To validate these experiments, the authors compared the results obtained by image cytometry with the traditional plating of yeast and bacterial cultures on PDA (Potato, Dextrose, and Agar) and MRS media, respectively. In their work, Williamson and co-authors obtained coefficients of variation values ranging from 0.67 to 1.64% (their data were presented in a Log10 scale) in the quantification of *S. cerevisiae* cells using the traditional plating technique, while our work showed an average coefficient of variation of 1.25% (transforming our results to a Log10 scale). For the quantification of *L. plantarum* cells, Williamson and collaborators presented coefficient of variation values from 0.31 to 1.16% in the same technique, whereas our work obtained a coefficient of variation of 0.75% in the Spread-plate plating technique, confirming that the values obtained in this work are similar to those existing and reported in the literature.

According to the main goal of this work, the following information was gathered for all evaluated techniques: operating principle, analysis time, detectable cell diameter range, simultaneous quantification capacity of yeast and bacterial cells, advantages, disadvantages, and estimated equipment cost and provided in Table 5 below. Based on Table 5, several important factors can be considered for choosing the method to be used for a specific application or study. Parameters such as analysis time, simultaneous quantification capacity, or estimated equipment cost, for example, can be decisive for choosing the technique, and the set of characteristics of each one should be carefully considered so that an informed decision, as much as possible, can be made according to the application’s objectives and expectations.

**Table 5.**
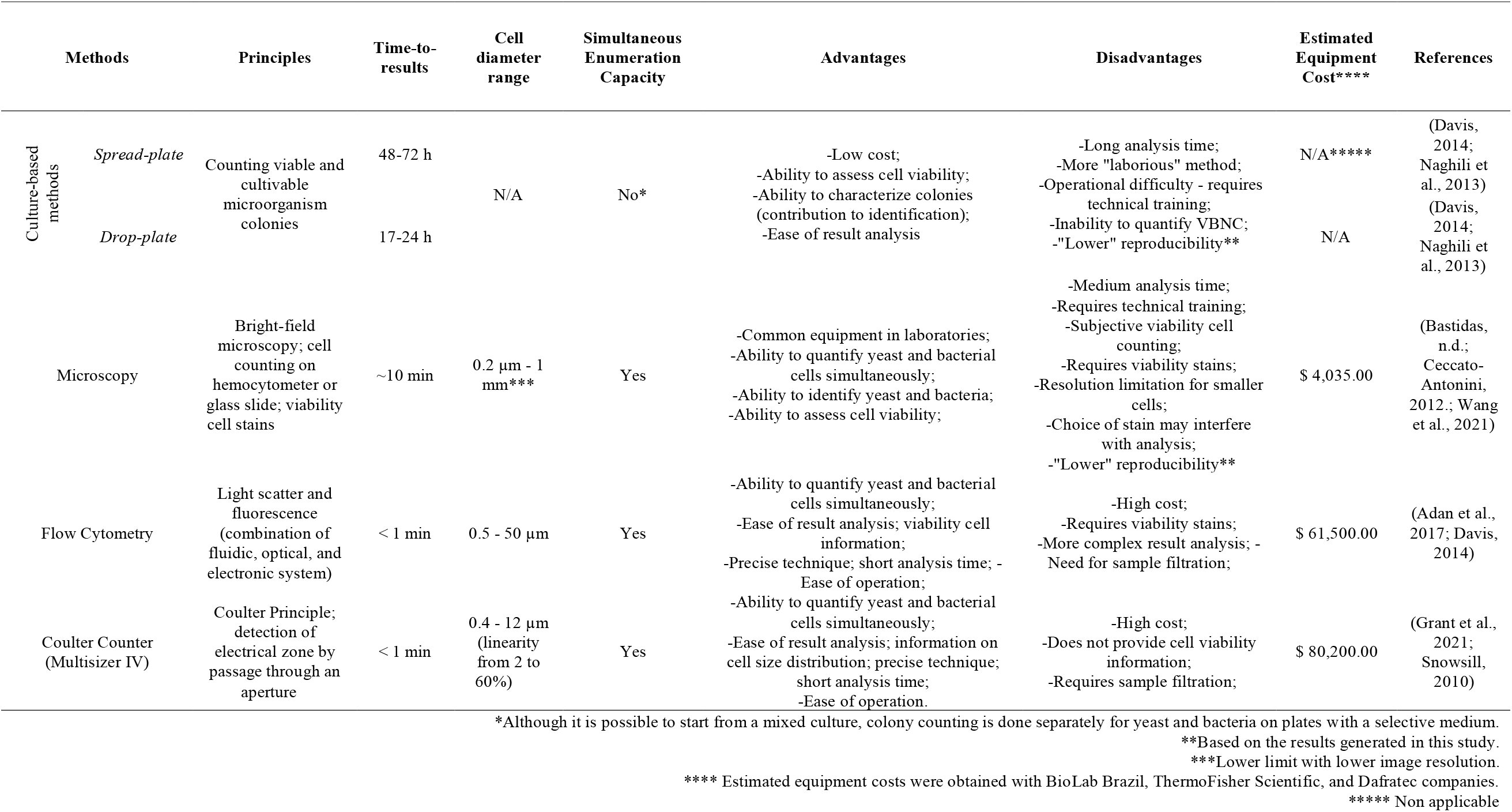
Key characteristics of each evaluated technique for simultaneous quantification of yeast and bacterial cells.

## Conclusion

In this work, we presented a comparison of the methods, in terms of time-to-results, cost of analysis and equipment, range of detectable cell/particle diameters, adequacy for simultaneous enumeration, and general pros and cons for yeast and bacterial simultaneous cell enumeration. The results obtained indicate that each method had its limitations, as well as its benefits in the simultaneous counting of yeast and bacterial cells and highlights how challenging is to pinpoint a single technique as ideal or better for all possible situations. For instance, while plate counts may underestimate microbial diversity and absolute cell concentration, flow cytometry, although efficient and rapid, may require expensive equipment and specialized training to analyze results. The choice of technique depends on various factors related to the development of academic research or research/application in industries, such as the budget available for obtaining equipment and reagents, qualified training, required result turnaround time, the accuracy and precision of obtained results, etc. Ideally, an integrated approach, combining complementary methods, might be used to gain a comprehensive understanding of the microbial community in the study. Additionally, it is crucial to consider the costs, ease of implementation, and specific limitations of each method in the decision-making process.

Additionally, it is worth noting that in this study, we did not start from real samples, for example, taken from fermentation vessels. The decision to begin the study with cell suspensions prepared from pure cultures of *S. cerevisiae* PE-2 and *L. plantarum* was guided by the choice to start with less complex cases, as industrial samples from biorefineries contain particulates that would increase the difficulty level of these analyses. Subsequent studies starting from real samples would greatly contribute to this discussion and deepen our knowledge and understanding of this environment.

## Acknowledgments

The authors are grateful to A. S. Lopes (UNICAMP), T. Granço (São Martinho), M. Lopes (Fermentec), C. Papini (Dafratec), J. Barreto (UNESP), and J. Gross (UNESP) for general inputs and discussions.

## Supplementary Material

Supplementary material is available online at JIMB (link).

