## Supplementary material for "Simultaneous enumeration of yeast and bacterial cells in the context of industrial bioprocesses"

**Supplementary file 1.** Python code applied for automated colony unit counting in platting methods.

```
import cv2 as cv

import numpy as np

import pandas as pd

import matplotlib.pyplot as plt

import matplotlib.image as img

from os import path, listdir, getcwd



# Function to find approx circles (yeasts)

def find_circles(img, ker=(5,5), edges_degree=40, tolerance=28, min_r=5, max_r=30):

  kernel = np.ones(ker,np.uint8)



  grad = cv.morphologyEx(img, cv.MORPH_GRADIENT, kernel)



  circles = cv.HoughCircles(grad, cv.HOUGH_GRADIENT, 1, 17, param1=edges_degree, param2=tolerance, minRadius=min_r, maxRadius=max_r)

  circles = np.uint16(np.around(circles))

  return circles



# draw contours of found yeasts

def circular(img, circles):

  for i in circles[0,:]:

      # draw the outer circle

      cv.circle(img,(i[0],i[1]),i[2],(255,0,0),2)

      # draw the center of the circle

      cv.circle(img,(i[0],i[1]),1,(0,75,75),3)



def find_yeasts_petri_camera(fileName):

if (path.isfile(fileName)):

        img_base = cv.imread(fileName)

        img = cv.cvtColor(img_base, cv.COLOR_BGR2GRAY)



        kernel = np.ones((5,5),np.uint8)

        gaus = cv.GaussianBlur(img,(5,5), 0)
```

```
		grad = cv.morphologyEx(gaus, cv.MORPH_GRADIENT, kernel)

		_, th = cv.threshold(grad, 23, 255, cv.THRESH_BINARY)


		cir = find_circles(th, ker=(5,5), edges_degree=40, tolerance=22, min_r=5, max_r=25)

		circular(img_base, cir)


else:

		print("Imagem nao encontrada")

		def find_yeasts_lamina(fileName):

if (path.isfile(fileName)):

		img_base = cv.imread(fileName)

		img = cv.cvtColor(img_base, cv.COLOR_BGR2GRAY)

		_, th1 = cv.threshold(img, 100, 255, cv.THRESH_BINARY_INV)

		cir = find_circles(th1, ker=(5,5), edges_degree=40, tolerance=28, min_r=5, max_r=30)

		circular(img_base, cir)

else:

		print("Imagem nao encontrada")


arq = input('Nome do arquivo: ')

tipo = int(input('\n1 - lamina\n2 - camera\n\nEscolha um numero: '))

print(tipo)

cwd = getcwd()

fileName = f'{cwd}/{arq}'


if (tipo == 1):

find_yeasts_lamina(fileName)

elif (tipo == 2):

find_yeasts_petri_camera(fileName)
```

**Table S1**. Results of the "Slide Method" and Neubauer Chamber yeast cell enumeration validation.

| | Technique | Unit | Rep1 | Rep2 | Rep3 | Rep4 | Mean | s.d | Coefficient of variation (%) |
|---|---|---|---|---|---|---|---|---|---|
| *S. cerevisiae (PE-2)* | *Neubauer Chamber* | cell/mL x10$^{-7}$ | 5.06 | 5.50 | 5.82 | 5.92 | **5.58** | *0.39* | 6.98 |
| | *"Slide Method"* | cell/mL x10$^{-7}$ | 4.00 | 4.44 | 6.60 | 5.99 | **5.26** | *1.24* | 23.57 |

**Table S2**. Table of ANOVA results and Tukey's t-test applied to the quantification of yeasts in mixed sample.

*ANOVAOneWay (04/10/2023 14:34:11)*

*Notes*

*Input Data*

*Descriptive Statistics*

| | Sample Size | Mean | Standard Deviation | SE of Mean |
|---|---|---|---|---|
| drop-plate | 5 | 7,8E7 | 9,08295E6 | 4,06202E6 |
| spread-plate | 5 | 4,16E7 | 9,09945E6 | 4,0694E6 |
| spread-plate automático | 5 | 4,2E7 | 7,74597E6 | 3,4641E6 |
| microscopia | 5 | 8,124E7 | 1,37546E7 | 6,15123E6 |
| multisizer 1:1 | 5 | 3,888E7 | 2,23428E6 | 999199,67974 |
| citometria 1:1 | 5 | 3,402E7 | 1,7225E6 | 770324,6069 |
| citometria 100:1 | 5 | 2,988E7 | 2,04377E6 | 914002,18818 |

*One Way ANOVA*

*Overall ANOVA*

| | DF | Sum of Squares | Mean Square | F Value | Prob>F |
|---|---|---|---|---|---|
| Model | 6 | 1,33779E16 | 2,22966E15 | 36,58394 | 5,39946E-12 |
| Error | 28 | 1,7065E15 | 6,09463E13 | | |
| Total | 34 | 1,50844E16 | | | |

Null Hypothesis: The means of all levels are equal.
Alternative Hypothesis: The means of one or more levels are different.
At the 0.05 level, the population means are significantly different.

*Fit Statistics*

| R-Square | Coeff Var | Root MSE | Data Mean |
|---|---|---|---|
| 0,88687 | 0,15811 | 7,80681E6 | 4,93743E7 |

*Means Comparisons*

*Tukey Test*

| | MeanDiff | SEM | q Value | Prob | Alpha | Sig | LCL | UCL |
|---|---|---|---|---|---|---|---|---|
| Spread-plate Drop-plate | -3,64E7 | 4,93746E6 | 10,42588 | 1,00382E-6 | 0,05 | 1 | -5,20623E7 | -2,07377E7 |
| Spread-plate auto Drop-plate | -3,6E7 | 4,93746E6 | 10,31131 | 1,2231E-6 | 0,05 | 1 | -5,16623E7 | -2,03377E7 |
| Spread-plate auto Spread-plate | 400000 | 4,93746E6 | 0,11457 | 1 | 0,05 | 0 | -1,52623E7 | 1,60623E7 |
| Microscopy Drop-plate | 3,24E6 | 4,93746E6 | 0,92802 | 0,99398 | 0,05 | 0 | -1,24223E7 | 1,89023E7 |
| Microscopy Spread-plate | 3,964E7 | 4,93746E6 | 11,3539 | 2,22348E-7 | 0,05 | 1 | 2,39777E7 | 5,53023E7 |
| Microscopy Spread-plate auto | 3,924E7 | 4,93746E6 | 11,23933 | 2,61695E-7 | 0,05 | 1 | 2,35777E7 | 5,49023E7 |
| Multisizer Drop-plate | -3,912E7 | 4,93746E6 | 11,20496 | 2,75256E-7 | 0,05 | 1 | -5,47823E7 | -2,34577E7 |
| Multisizer Spread-plate | -2,72E6 | 4,93746E6 | 0,77908 | 0,99768 | 0,05 | 0 | -1,83823E7 | 1,29423E7 |
| Multisizer Spread-plate auto | -3,12E6 | 4,93746E6 | 0,89365 | 0,99509 | 0,05 | 0 | -1,87823E7 | 1,25423E7 |
| Multisizer Microscopy | -4,236E7 | 4,93746E6 | 12,13298 | 8,16055E-8 | 0,05 | 1 | -5,80223E7 | -2,66977E7 |
| Cytometry 1:1 Drop-plate | -4,398E7 | 4,93746E6 | 12,59698 | 5,11675E-8 | 0,05 | 1 | -5,96423E7 | -2,83177E7 |
| Cytometry 1:1 Spread-plate | -7,58E6 | 4,93746E6 | 2,1711 | 0,72206 | 0,05 | 0 | -2,32423E7 | 8,08229E6 |
| Cytometry 1:1 Spread-plate auto | -7,98E6 | 4,93746E6 | 2,28567 | 0,67349 | 0,05 | 0 | -2,36423E7 | 7,68229E6 |
| Cytometry 1:1 Microscopy | -4,722E7 | 4,93746E6 | 13,525 | 4,67012E-8 | 0,05 | 1 | -6,28823E7 | -3,15577E7 |
| Cytometry 1:1 Multisizer | -4,86E6 | 4,93746E6 | 1,39203 | 0,95333 | 0,05 | 0 | -2,05223E7 | 1,08023E7 |
| Cytometry 100:1 Drop-plate | -4,812E7 | 4,93746E6 | 13,78279 | 4,38143E-8 | 0,05 | 1 | -6,37823E7 | -3,24577E7 |
| Cytometry 100:1 Spread-plate | -1,172E7 | 4,93746E6 | 3,3569 | 0,24668 | 0,05 | 0 | -2,73823E7 | 3,94229E6 |
| Cytometry 100:1 Spread-plate auto | -1,212E7 | 4,93746E6 | 3,47147 | 0,21427 | 0,05 | 0 | -2,77823E7 | 3,54229E6 |
| Cytometry 100:1 Microscopy | -5,136E7 | 4,93746E6 | 14,7108 | 3,71256E-8 | 0,05 | 1 | -6,70223E7 | -3,56977E7 |
| Cytometry 100:1 Multisizer | -9E6 | 4,93746E6 | 2,57783 | 0,54462 | 0,05 | 0 | -2,46623E7 | 6,66229E6 |
| Cytometry 100:1 Cytometry 1:1 | -4,14E6 | 4,93746E6 | 1,1858 | 0,97851 | 0,05 | 0 | -1,98023E7 | 1,15223E7 |

Sig equals 1 indicates that the means difference is significant at the 0,05 level.
Sig equals 0 indicates that the means difference is not significant at the 0,05 level.

**Table S3.** Table of ANOVA results and Tukey's t-test applied to the quantification of bacteria in mixed sample.

*ANOVAOneWay (04/10/2023 14:52:42)*

*Notes*

*Input Data*

*Descriptive Statistics*

| | Sample Size | Mean | Standard Deviation | SE of Mean |
|---|---|---|---|---|
| drop-plate | 5 | 2,17E9 | 1,39642E8 | 6,245E7 |
| spread-plate | 5 | 2,192E9 | 3,33946E8 | 1,49345E8 |
| spread-plate automático | 5 | 2,004E9 | 2,6397E8 | 1,18051E8 |
| microscopia | 5 | 2,404E9 | 7,03086E8 | 3,1443E8 |
| multisizer 1:1 | 5 | 4,868E9 | 4,42854E8 | 1,9805E8 |
| citometria 1:1 | 5 | 1,474E9 | 1,42934E8 | 6,39218E7 |
| citometria 100:1 | 5 | 3,156E9 | 2,20624E9 | 9,86659E8 |

*One Way ANOVA*

*Overall ANOVA*

| | DF | Sum of Squares | Mean Square | F Value | Prob>F |
|---|---|---|---|---|---|
| Model | 6 | 3,73259E19 | 6,22098E18 | 7,53528 | 7,11782E-5 |
| Error | 28 | 2,31162E19 | 8,2558E17 | | |
| Total | 34 | 6,04421E19 | | | |

Null Hypothesis: The means of all levels are equal.
Alternative Hypothesis: The means of one or more levels are different.
At the 0.05 level, the population means are significantly different.

*Fit Statistics*

| R-Square | Coeff Var | Root MSE | Data Mean |
|---|---|---|---|
| 0,61755 | 0,34817 | 9,08614E8 | 2,60971E9 |

*Means Comparisons*

*Tukey Test*

| | MeanDiff | SEM | q Value | Prob | Alpha | Sig | LCL | UCL |
|---|---|---|---|---|---|---|---|---|
| Spread-plate Drop-plate | 2,2E7 | 5,74658E8 | 0,05414 | 1 | 0,05 | 0 | -1,80089E9 | 1,84489E9 |
| Spread-plate auto Drop-plate | -1,66E8 | 5,74658E8 | 0,40852 | 0,99994 | 0,05 | 0 | -1,98889E9 | 1,65689E9 |
| Spread-plate auto Spread-plate | -1,88E8 | 5,74658E8 | 0,46266 | 0,99988 | 0,05 | 0 | -2,01089E9 | 1,63489E9 |
| Microscopy Drop-plate | 2,34E8 | 5,74658E8 | 0,57587 | 0,99958 | 0,05 | 0 | -1,58889E9 | 2,05689E9 |
| Microscopy Spread-plate | 2,12E8 | 5,74658E8 | 0,52172 | 0,99976 | 0,05 | 0 | -1,61089E9 | 2,03489E9 |
| Microscopy Spread-plate auto | 4E8 | 5,74658E8 | 0,98439 | 0,99176 | 0,05 | 0 | -1,42289E9 | 2,22289E9 |
| Multisizer Drop-plate | 2,698E9 | 5,74658E8 | 6,63968 | 0,00112 | 0,05 | 1 | 8,75107E8 | 4,52089E9 |
| Multisizer Spread-plate | 2,676E9 | 5,74658E8 | 6,58554 | 0,00124 | 0,05 | 1 | 8,53107E8 | 4,49889E9 |
| Multisizer Spread-plate auto | 2,864E9 | 5,74658E8 | 7,0482 | 5,19546E-4 | 0,05 | 1 | 1,04111E9 | 4,68689E9 |
| Multisizer Microscopy | 2,464E9 | 5,74658E8 | 6,06382 | 0,00325 | 0,05 | 1 | 6,41107E8 | 4,28689E9 |
| Cytometry 1:1 Drop-plate | -6,96E8 | 5,74658E8 | 1,71283 | 0,88395 | 0,05 | 0 | -2,51889E9 | 1,12689E9 |
| Cytometry 1:1 Spread-plate | -7,18E8 | 5,74658E8 | 1,76697 | 0,86838 | 0,05 | 0 | -2,54089E9 | 1,10489E9 |
| Cytometry 1:1 Spread-plate auto | -5,3E8 | 5,74658E8 | 1,30431 | 0,96571 | 0,05 | 0 | -2,35289E9 | 1,29289E9 |
| Cytometry 1:1 Microscopy | -9,3E8 | 5,74658E8 | 2,2887 | 0,67218 | 0,05 | 0 | -2,75289E9 | 8,92893E8 |
| Cytometry 1:1 Multisizer | -3,394E9 | 5,74658E8 | 8,35251 | 4,4349E-5 | 0,05 | 1 | -5,21689E9 | -1,57111E9 |
| Cytometry 100:1 Drop-plate | 9,86E8 | 5,74658E8 | 2,42651 | 0,61172 | 0,05 | 0 | -8,36893E8 | 2,80889E9 |
| Cytometry 100:1 Spread-plate | 9,64E8 | 5,74658E8 | 2,37237 | 0,63564 | 0,05 | 0 | -8,58893E8 | 2,78689E9 |
| Cytometry 100:1 Spread-plate auto | 1,152E9 | 5,74658E8 | 2,83503 | 0,43402 | 0,05 | 0 | -6,70893E8 | 2,97489E9 |
| Cytometry 100:1 Microscopy | 7,52E8 | 5,74658E8 | 1,85065 | 0,84224 | 0,05 | 0 | -1,07089E9 | 2,57489E9 |
| Cytometry 100:1 Multisizer | -1,712E9 | 5,74658E8 | 4,21317 | 0,07634 | 0,05 | 0 | -3,53489E9 | 1,10893E8 |
| Cytometry 100:1 Cytometry 1:1 | 1,682E9 | 5,74658E8 | 4,13934 | 0,0853 | 0,05 | 0 | -1,40893E8 | 3,50489E9 |

Sig equals 1 indicates that the means difference is significant at the 0,05 level.
Sig equals 0 indicates that the means difference is not significant at the 0,05 level.

**Table S4.** Values of cell viability replicates obtained for yeasts.

| Yeast cell viability (%) | | | | | | | |
|---|---|---|---|---|---|---|---|
| | Rep 1 | Rep 2 | Rep 3 | Rep 4 | Rep 5 | Mean (%) | S.D (%) |
| Bright field microscopy (Erythrosine B) | 93.0 | 93.1 | 94.3 | 88.5 | 89.0 | 91.6 | 2.9 |
| Flow cytometry (Propidium Iodide - ratio 1:1) | 90.0 | 90.0 | 92.0 | 90.0 | 90.0 | 90.4 | 0.9 |

**Table S5**. Values of cell viability replicates obtained for bacterial cells.

| Bacterial cell viability (%) | | | | | | | |
|---|---|---|---|---|---|---|---|
| | Rep 1 | Rep 2 | Rep 3 | Rep 4 | Rep 5 | Mean (%) | S.D (%) |
| Bright field microscopy (Erythrosine B) | 89 | 92 | 91 | 86 | 86 | 88.8 | 3.1 |
| Flow cytometry (Propidium Iodide - ratio 1:1) | 99 | 98 | 99 | 100 | 99 | 99 | 0.71 |

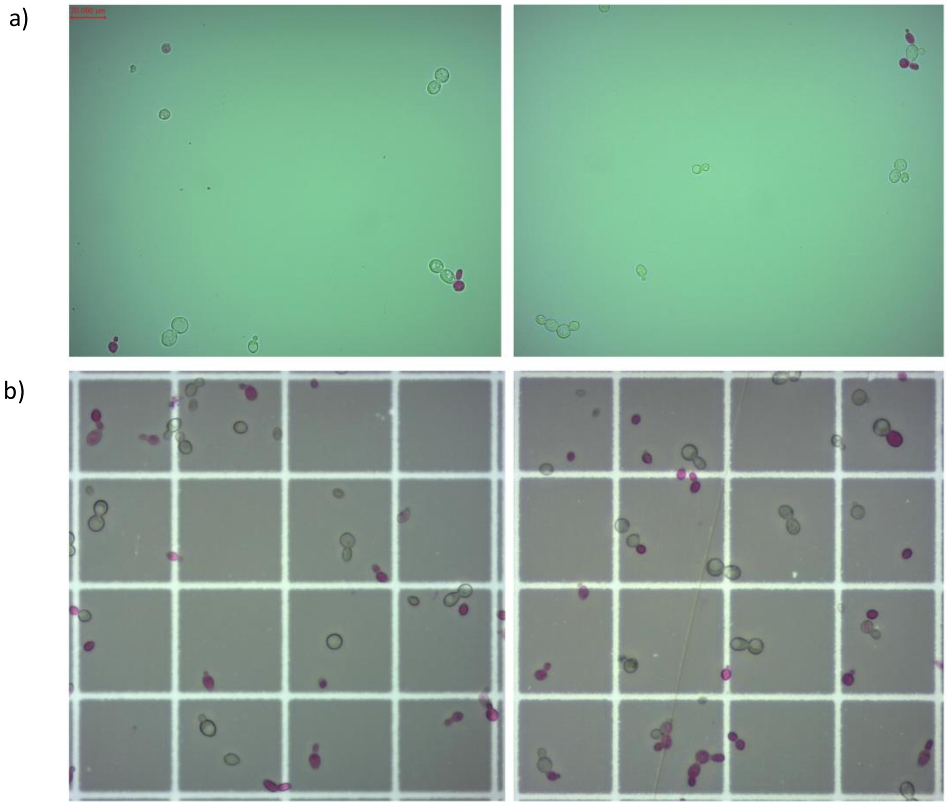


**Figure S1.** Microscopy images of the "Slide Method" and Neubauer Chamber yeast cell enumeration validation. A) "Slide Method", and b) Neubauer Chamber

a)

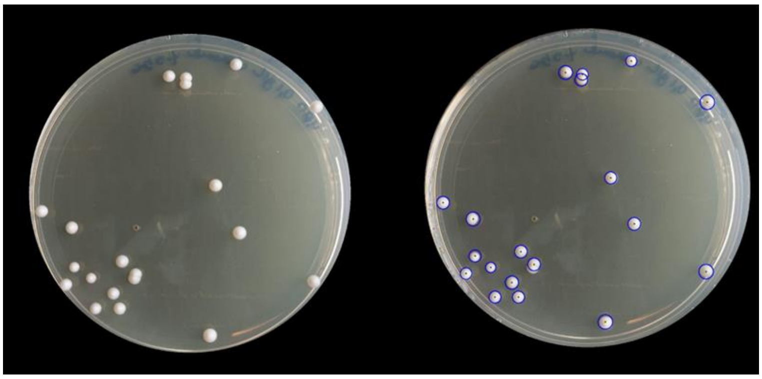

b)

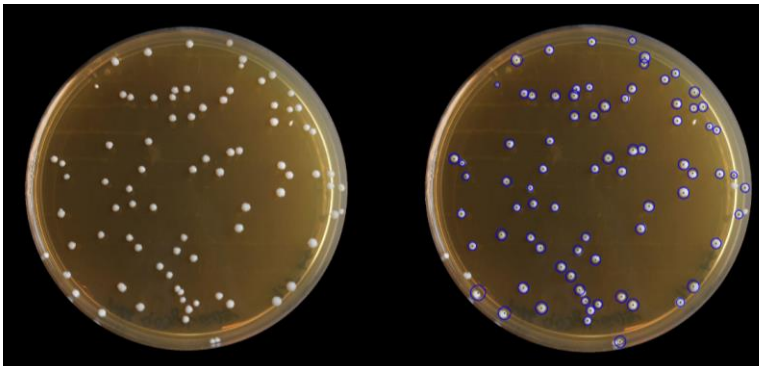

**Figure S2**. Plate containing YPD-agar with chloramphenicol (100 μg/mL) (Figure S2a) and a plate containing MRS-agar with cycloheximide (50 μg/mL) (Figure S2b), on which the mixed cell suspension of yeast and bacterial cells (in a cell ratio of 1:1) was spread and incubated. After the appropriate incubation period, manual counting was performed (Figure S2, left panel), and a photographic record of each plate was taken for subsequent automatic counting (Figure S2, right panel). The overlaid blue circles on each colony (image on the right) represent the area of automatic identification and counting.

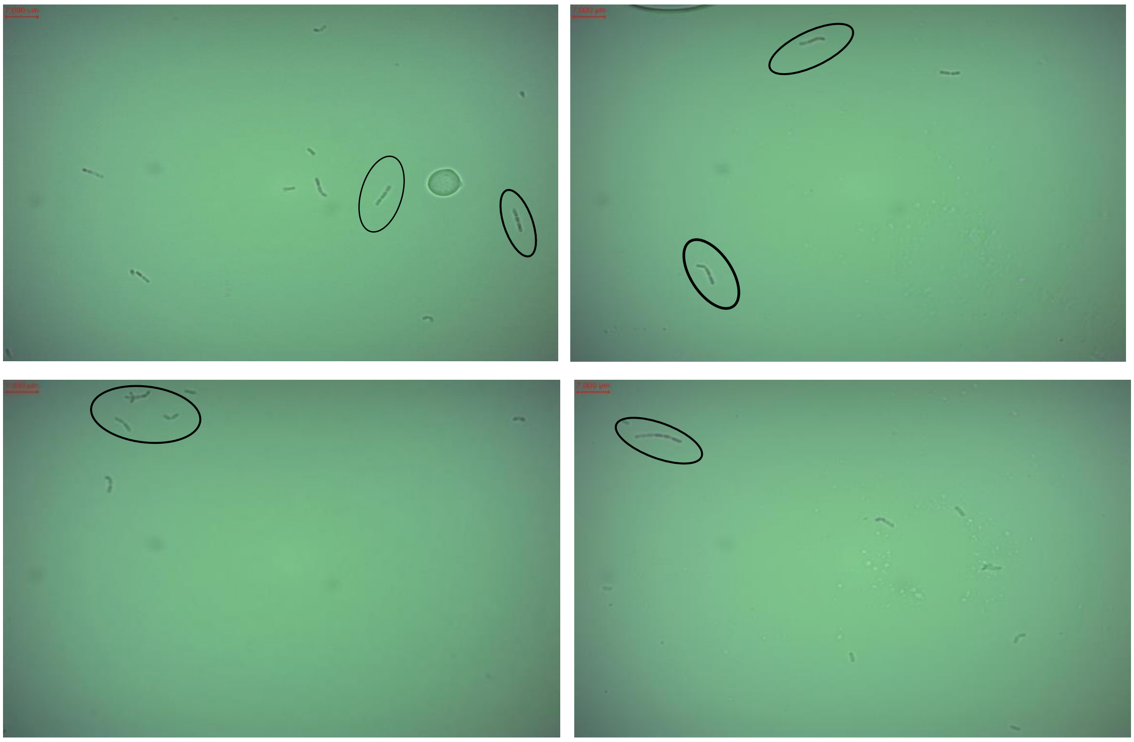

**Figure S3**. Images of the mixed suspension of *S. cerevisiae* and *L. plantarum* cells by the adapted bright-field microscopy, with a 1000x magnification and immersion oil. The *L. plantarum* cells in "chains" are highlighted by black ellipses.
